# Compound-specific, concentration-independent biophysical properties of sodium channel inhibitor mechanism of action

**DOI:** 10.1101/2021.07.05.451191

**Authors:** Krisztina Pesti, Mátyás C. Földi, Katalin Zboray, Adam V. Toth, Peter Lukacs, Arpad Mike

## Abstract

We have developed an automated patch-clamp protocol that allows high information content screening of sodium channel inhibitor compounds. We have observed that individual compounds had their specific signature patterns of inhibition, which were manifested irrespective of the concentration. Our aim in this study was to quantify these properties. Primary biophysical data, such as onset rate, the shift of the half inactivation voltage, or the delay of recovery from inactivation, are concentration-dependent. We wanted to derive compound-specific properties, therefore, we had to neutralize the effect of concentration. This study describes how this is done, and shows how compound-specific properties reflect the mechanism of action, including binding dynamics, cooperativity, and interaction with the membrane phase. We illustrate the method using four well-known sodium channel inhibitor compounds, riluzole, lidocaine, benzocaine, and bupivacaine. Compound-specific biophysical properties may also serve as a basis for deriving parameters for kinetic modeling of drug action. We discuss how knowledge about the mechanism of action may help to predict the frequency-dependence of individual compounds, as well as their potential persistent current component selectivity. The analysis method described in this study, together with the experimental protocol described in the accompanying paper, allows screening for inhibitor compounds with specific kinetic properties, or with specific mechanisms of inhibition.

## Introduction

*In silico* prediction of drug effects can save a tremendous amount of time and resources, and can accelerate drug discovery. To predict the therapeutic action of sodium channel inhibitors, an elementary knowledge about the mechanism of action is essential. These drugs show state-dependent accessibility and affinity to binding sites and can have radically different binding/unbinding kinetics. These properties determine their effect, as they dynamically bind and unbind depending on the activity pattern of individual cells.

*In silico* prediction is especially useful for multi-target drugs, where targets are not independent of each other but interact in a complex way. Voltage-gated ion channels both affect and are affected by the membrane potential. Similarly, they both affect and are affected by intracellular ion concentrations. This creates an intricate network of interactions, the outcome of which is difficult to predict without modeling. Ion channels also happen to be the most promiscuous drug targets, therefore multi-target effects among therapeutic drugs are much rather the rule, than the exception^1–4^. Predicting the effect of a specific drug – either to achieve therapeutic effects or to avoid adverse effects – in most cases requires considering its interaction with several ion channel and other targets, together with the complex network of interactions between the targets themselves. The best-studied example for this is the torsadogenic effect of certain compounds in the human heart, where *in silico* modeling of multi-target effects is now a generally accepted directive^5^. This initially has been done by determining the IC_50_ value of a specific drug to all relevant ion channel targets, from which the inhibited fraction of specific ion channels can be determined at specific drug concentrations. These conductances then were reduced according to the inhibited fraction in the simulations. However, as it has been first shown by Di Veroli *et al*.^6,7^, the predicted effect can be seriously underestimated if one does not consider the mechanism of action; most importantly the dynamics of perpetual state-dependent binding/unbinding. Indeed, simulated compounds at their IC_50_ concentration could have widely different effects on the action potential duration depending on their state preference and binding kinetics^8^. Some of the more recent, improved models, therefore, include Markov models of hERG channels, where state-dependence and binding/unbinding dynamics is also simulated^8–10^. The same approach should be applied for modeling drug effects on sodium channels, not only in the context of cardiac safety pharmacology, but in predicting therapeutic efficacy for all hyperexcitability-related conditions including neuromuscular disorders, pain syndromes, epilepsies, and cardiac arrhythmias. A dependable model should include several processes, such as aqueous phase – membrane phase partitioning, state-dependent access (as described by the guarded receptor hypothesis^11^), and state-dependent affinity, which is inevitably linked with allosteric modulation of channel gating (as described by the modulated receptor hypothesis^12^). The mechanism of inhibition for most sodium channel inhibitor drugs is not known in sufficient detail to allow the construction of adequate models, and a comprehensive analysis of mechanisms for a reasonable number of sodium channel inhibitors is still lacking. In the accompanying study, we aimed to develop a protocol by which an initial assessment of these processes can be completed with reasonably high throughput. In this study, our aim was to characterize the mechanism of action of individual drugs, not a specific concentration of a specific drug. For example, one may observe that the onset of effect for compound “A” is faster than for compound “B”, or that compound “B” delays recovery more effectively than compound “A”. Does this tell anything about their specific mechanisms of action? Not necessarily. It is possible that if we increase the concentration of compound “B”, the onset will be just as fast as that of compound “A”. It is also possible that if we increase the concentration of compound “A”, it will delay recovery just as effectively as compound “B”. We aimed to find compound-specific (and concentration-independent) properties of inhibition, and it turned out that each compound did have such properties, not only resting and inactivated state affinities (*K_R_* and *K_I_*), but more importantly the kinetics of approaching *K_R_* upon hyperpolarization, and approaching *K_I_* upon depolarization were also such compound-specific properties.

## Methods

### Cell culture and Automated patch-clamp electrophysiology

Cell culture and electrophysiology were done as described in the accompanying paper^13^. The recombinant rNaV1.4 channel-expressing cell line was generated as described before^14^. Transfected HEK 293 cells were maintained in Dulbecco’s Modified Eagle Medium, high glucose supplemented with 10% v/v fetal bovine serum, 100 U/ml of penicillin/streptomycin, and 0.4 mg/mL Geneticin (Life Technologies, Carlsbad, CA). For experiments, cells were dissociated from culture dishes with Accutase (Corning), shaken in serum-free medium for 60 minutes at room temperature, then centrifuged, and resuspended into the extracellular solution to a concentration of 5×10^6^ cells/mL. Ensemble voltage-clamp recordings were performed on an IonFlux Mercury instrument (Fluxion Biosciences). The composition of solutions (in mM) was: Intracellular solution: 50 CsCl, 10 NaCl, 60 CsF, 20 EGTA, 10 HEPES; pH 7.2 (adjusted with 1 M CsOH). Extracellular solution: 140 NaCl, 4 KCl, 1 MgCl_2_, 2 CaCl_2_, 5 D-Glucose and 10 HEPES; pH 7.4 (adjusted with 1 M NaOH). The osmolality of intra- and extracellular solutions was set to ~320 and ~330 mOsm, respectively. Data were sampled at 20 kHz, and filtered at 10 kHz. Experiments were carried out at room temperature. The 384-well IonFlux microfluidic ensemble plates are divided into four “zones”, typically each zone was used for a separate experiment (one particular set of compounds on one particular cell line). Each zone consists of 8 separate sections, which are distinct functional units, containing one well for the cell suspension, one well for the waste, two cell “traps” (intracellular solution-filled wells under negative pressure to establish high resistance seals and then whole-cell configuration), and eight compound wells. We kept one compound well for cell-free extracellular solution and typically used the remaining 7 compound plates for 2 different compounds one of them in 3, the other in 4 different concentrations. From the 16 cell ensembles of each zone, we chose n = 6 ensembles for analysis, based on the stability of amplitude and seal resistance throughout the experiment.

We used a complex 17-pulse voltage protocol (Fig. 1A), described in detail in the accompanying paper^13^, which allowed the assessment of gating kinetics and gating equilibrium in the absence and the presence of inhibitor/modulator compounds. The protocol investigated the effect of different durations of depolarizations “state-dependent onset” (**SDO**), the effect of different durations of hyperpolarizations “recovery from inactivation” (**RFI**), and the resting-inactivated equilibrium at different membrane potentials “steady-state inactivation” (**SSI**), similarly to protocols we used in previous studies^14,15^.

**Fig. 1.**
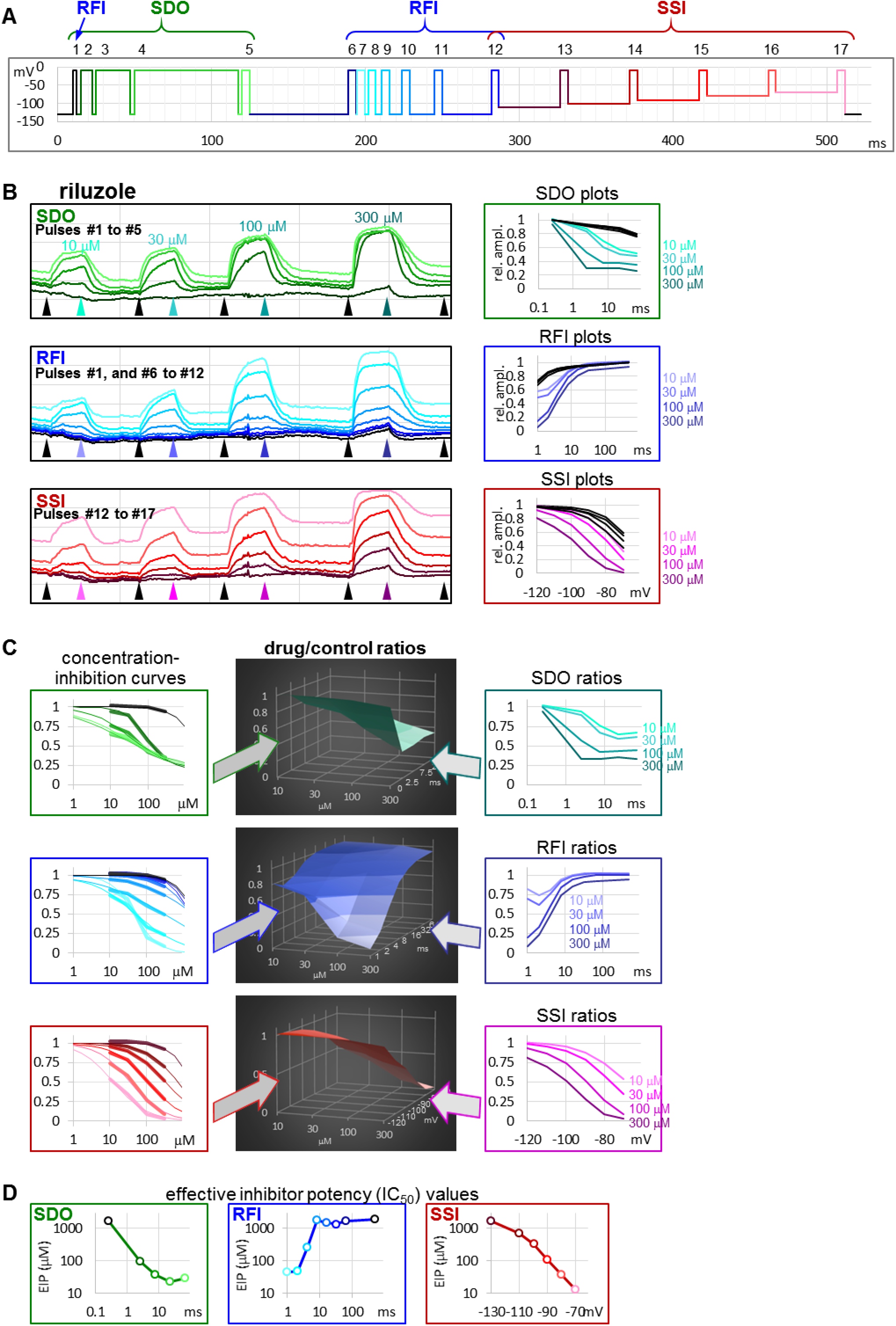
The sequence of analysis to obtain effective inhibitor potency plots. The process is illustrated in an example of an experiment, where four concentrations of riluzole were applied to a cell ensemble. **(A)** Schematic picture of the voltage protocol, which was repeated at 1 Hz frequency. The color of pulses match the color of amplitude plots in panel B, the color of concentration-inhibition curves in panel C, left column, and the color of open circles in EIP plots in panel D. The number of pulses, and the three main sections of the protocol: **SDO**, **RFI**, and **SSI**, are indicated. **(B) Left column:** Amplitude plots throughout the four riluzole applications, for the three main sections. Grid size: 1 nA (vertical), 100 s (horizontal). Arrowheads show the time points, from where data were collected to construct plots in the right column. **Right column: SDO**, **RFI**, and **SSI** plots for four different concentrations of riluzole, and four sets of control data before drug application. The color of curves match the color of arrowheads in the left column. **(C) Middle column:** 3D plots of drug/control ratios for each of the four concentrations (front axis), and for each of the 17 pulses (three different right-side axes at the three main sections). **Left column:** Concentration-inhibition plots for each of the 17 pulses (Projection of middle column 3D plots to the front plane). Thick lines show experimental data, thin lines show fits of the Hill equation to the data. **Right column:** Drug/control ratios shown on the same abscissae as **SDO**, **RFI**, and **SSI** plots in panel B, left column (Projection of middle column 3D plots to the right side plane). **(D)** Effective inhibitor potency plots show how EIP of riluzole was found to change depending on conditioning pulse duration (**SDO**), interpulse interval (**RFI**), and holding potential (**SSI**)

### Data analysis

In addition to automatic fitting, described in the accompanying paper^13^, in this study we used a different approach to process **SDO**, **RFI**, and **SSI** data. This analysis focused on deriving compound-specific but concentration-independent descriptors of the mechanism of action, which can lead to a mode thorough understanding of the sub-processes involved in drug action, and which can later serve as a basis for constructing kinetic models for the simulation of drug-specific effects.

Inhibition of 1^st^ and 17^th^ pulse-evoked current amplitudes were used to obtain estimates of resting-state-, and inactivated-state-affinities (*K_R_* and *K_I_*). We constructed concentration-inhibition curves for both the 1^st^ and 17^th^ pulse-evoked currents, and fitted them with the Hill equation:

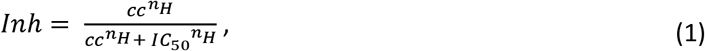

where *Inh* is the inhibited fraction of the current, *cc* is the drug concentration, and *n_H_* is the Hill coefficient. *K_R_* was approximated with the *IC_50_* value for 1^st^ pulse-evoked currents. For the calculation of *K_I_*, we used the equation from Bean et al.^16^:

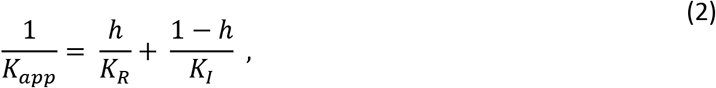

where *K_app_* was the *IC_50_* for the 17^th^ pulse, and (*1-h*) was the ratio of 17^th^/1^st^ pulse evoked current amplitudes (from the average of the last 5 s before drug application; see pink traces (pulse #17) and black traces (pulse #1) in Fig. 1B). *K_R_* and *K_I_* were the extreme values between which the potency of inhibition continuously fluctuated, depending on the voltage protocol. A detailed description of the analysis of dynamic changes in potency is found below in Results.

## Results

In the accompanying paper^13^ we describe how different degrees of inhibition observed in the 17-pulse protocol can be interpreted as revealing the state-dependent onset (**SDO**; how the inhibition depended on the length of depolarizations), the recovery from inactivation (**RFI**; how the inhibition depended on the length of hyperpolarizations), and steady-state inactivation (**SSI**; how the inhibition depended on the membrane potential). We have distinguished “macro-dynamics”, which was the onset/offset upon drug perfusion and removal, and occurred on the second-timescale; and “micro-dynamics”, which was the onset/offset upon conformational transitions of the channel population, and occurred on the millisecond timescale. In this study we focus on micro-dynamics. We first explain the sequence of analysis on the example of riluzole (in four different concentrations) using data from a single cell ensemble. Then we show the same analysis on examples for three additional compounds in order to demonstrate how similar compounds may radically differ in their micro-dynamics. Finally, we show how these compound-specific biophysical properties may affect their action at actively firing excitable cells.

### Sequence of analysis, and the concept of “effective inhibitor potency”

The 17-pulse protocol is shown in Fig. 1A; it was repeated at each second (1 Hz). For the sake of clarity, we show the sequence of analysis on a single measurement (the ensemble of 20 simultaneously recorded cells) for different concentrations of riluzole. Peak amplitude plots for all 17 pulse-evoked currents are shown in Fig. 1B, left panel; peak amplitudes were arranged into three separate groups as shown in Fig. 1A, this allowed us to construct **SDO**, **RFI**, and **SSI** plots. In this procedure of analysis, we did not use every sweep of the experiment, only constructed one set of plots right before the start of each drug perfusion period (controls), as well as one set of plots at the end of each drug perfusion period (in this case 10, 30, 100, and 300 μM riluzole), as shown by the arrowheads in Fig. 1B. The constructed plots are shown in Fig. 1B, right panel.

It is obvious that the same concentration of riluzole differently affected currents evoked by different depolarizations. For example, the effect of 100 μM riluzole fluctuated between almost full inhibition (pulses #7 and #17) and no inhibition (pulses #1, #6, and #12). We termed this process “micro-dynamics”, to distinguish from the onset and offset of drug effects upon perfusion and washout of riluzole (“macro-dynamics”), which occurred on a ~1000-fold slower time scale (Fig. 1B, Table 1). For a more detailed discussion of micro- and macro-dynamics see the accompanying paper^13^. The term “apparent affinity” has been used to reflect different degrees of inhibition at different membrane potentials^17,18^. We have previously extended the use of this term to non-equilibrium conditions (such as during the course of recovery from inactivation)^15^, however, for discussing non-equilibrium conditions we consider it better to introduce the term “effective inhibitory potency” (EIP). The term “affinity” in itself conveys that the effect is the consequence of a binding/unbinding equilibrium. During the onset of inhibition, or recovery from inhibition, however, there is no equilibrium, and there is a complex combination of different processes beyond simple binding/unbinding, such as modulation of gating, access into/egress from the central cavity, aqueous phase-membrane partitioning, deprotonation/protonation, etc. The term “potency”, on the other hand, makes no reference to the mechanism, it simply expresses what fraction of the channel population can a certain concentration of a certain compound inhibit at a certain point in time. EIP, just like affinity, can be quantified by constructing concentration-response curves and determining the IC_50_ values. We found that the EIP kept changing dynamically, it increased (*i.e.,* IC_50_ values decreased) with longer conditioning pulses (**SDO**), and decreased with longer interpulse intervals (**RFI**). To calculate EIP for different conditions, we calculated drug-treatment/control ratios for all 17 pulses, and for all drug concentrations: we divided the amplitude of each 17 evoked currents in the presence of all four concentrations of riluzole by the corresponding control amplitude. The results are shown in Fig. 1C. All three columns show the exact same set of amplitude ratios. The middle column of 3D plots shows drug-treatment/control ratios both as a function of riluzole concentration, and as a function of conditioning pulse duration (**SDO**, upper row), interpulse interval (**RFI**, middle row), and membrane potential (**SSI**, lower row). The right column illustrates the projection of the 3D plot to the right side plane (as seen from the right side, shown by the teal, indigo, and purple arrows). These plots are rather similar to the ones shown in the right column of Fig. 1B, but current amplitudes evoked by each pulse have been normalized, each to its own control. The left column (thick lines) illustrates the projection of the 3D plot to the front plane (as seen from the front; shown by the green, blue, and red arrows), thus forming 17 concentration-response plots. Note that all 17 concentration-response curves were different. From the fitted Hill equations (thin lines) the IC_50_ values were determined and plotted against conditioning pulse duration (**SDO**), interpulse interval (**RFI**), or membrane potential (**SSI**)(Fig. 1D). Hill coefficients (*n_H_*) ranged between 0.5 and 2.0; some of the reasons why Hill coefficients may have diverged from unity will be discussed below. EIP plots for the **SDO** and **RFI** sections of the protocol show the major parameters of micro-dynamics for individual compounds: how fast its effect develops, and how fast it is terminated, depending on the conformational dynamics of the channel protein. We can observe that riluzole showed fast micro-dynamics: essentially both the onset and the offset of its effect were complete within ~10 ms. Micro-dynamics data from n = 6 cell ensembles will be shown below in Fig. 5A.

**Table 1.**
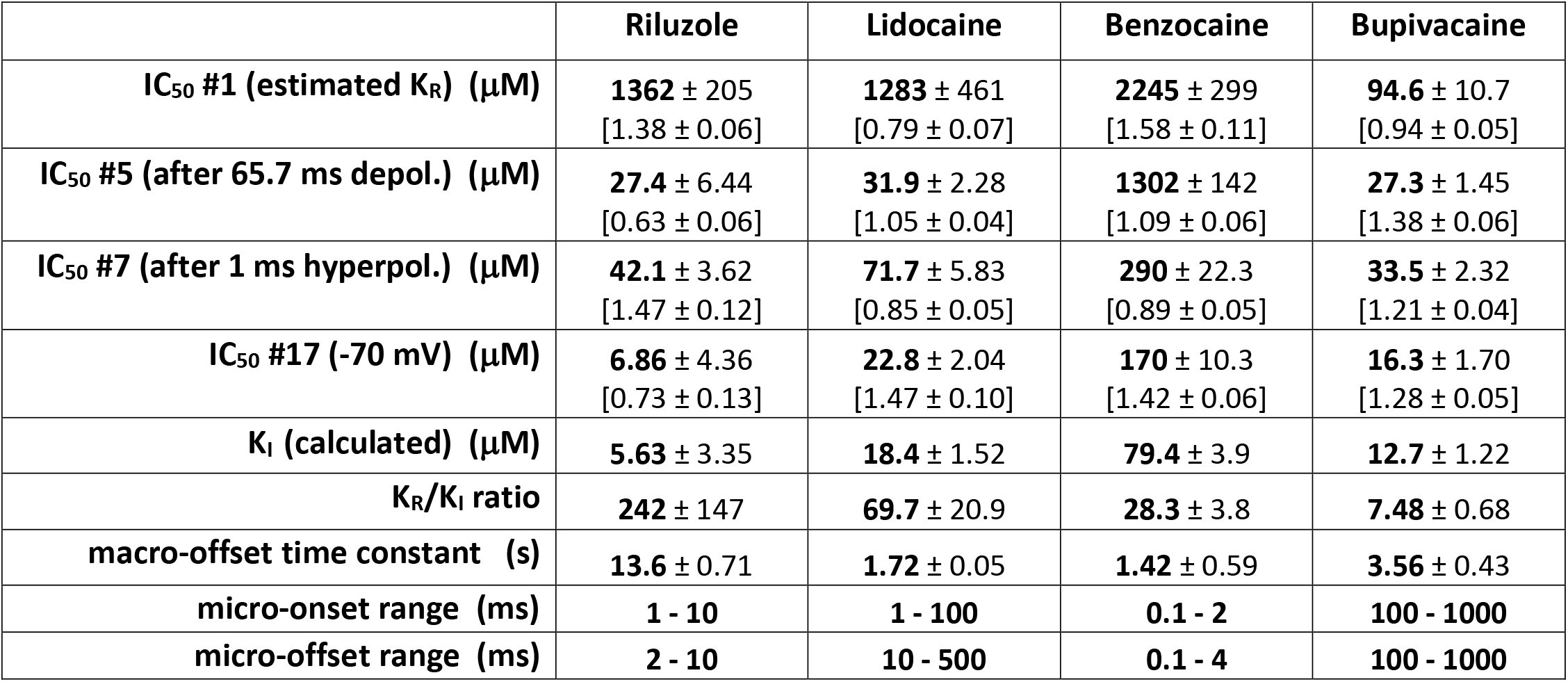
Main parameters of inhibition by four compounds. K_I_ was calculated as described in Methods. Micro-association and micro-dissociation could not be fit with a single exponential, therefore we give the range within which the effective inhibitor potency was observed to change.

**Fig. 2.**
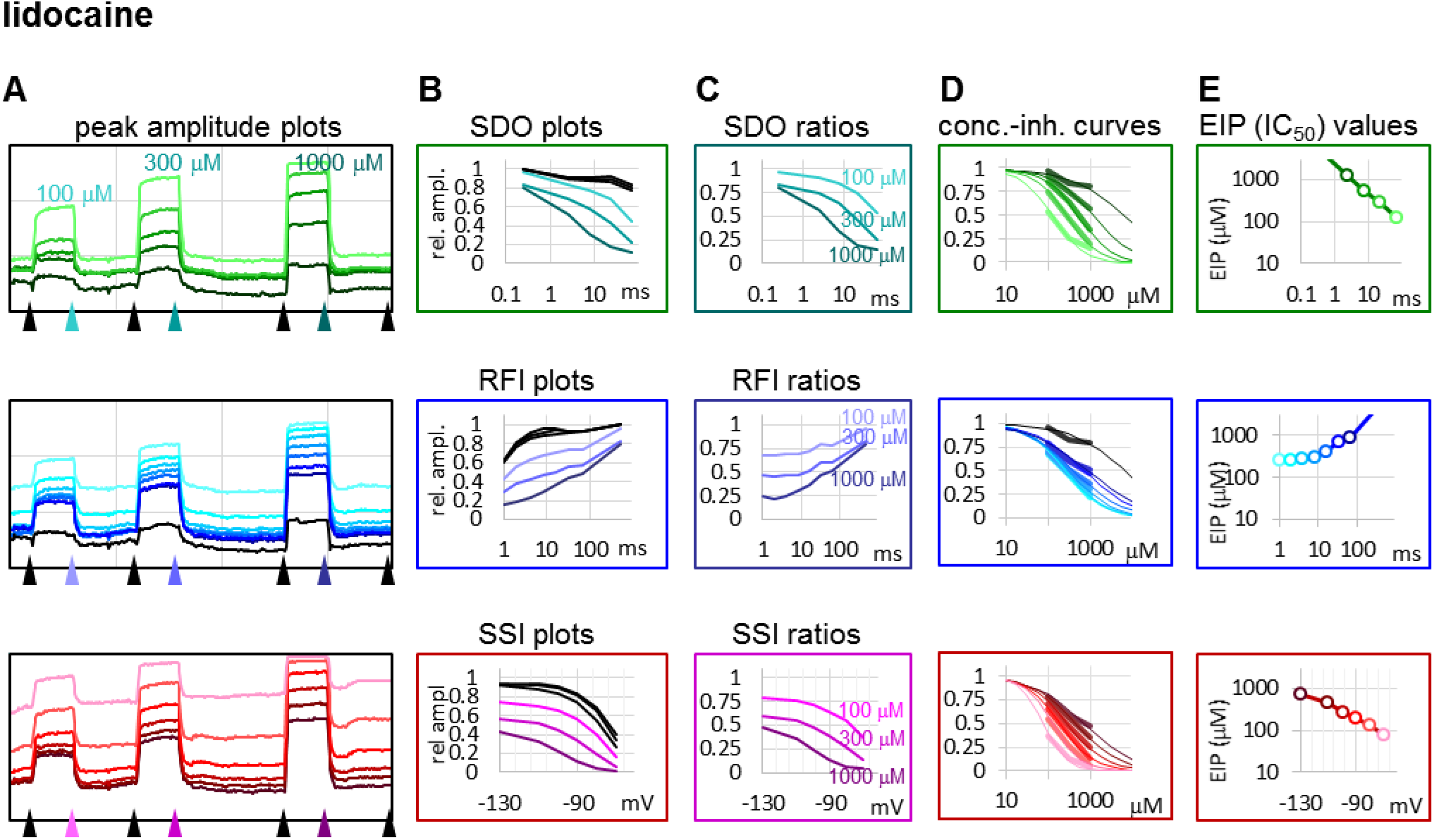
Construction of EIP plots for lidocaine. The figure illustrates how EIP plots are constructed on the example of a single measurement of the effect of three concentrations of lidocaine. **(A)** Peak amplitudes of currents evoked by the 17-pulse protocol were grouped into three sections: **SDO** (upper row), **RFI** (middle row), and **SSI** (lower row). **(B)** Amplitudes in control sections and during lidocaine perfusion were plotted against conditioning pulse duration (**SDO**), interpulse interval duration (**RFI**), and membrane potential (**SSI**). **(C)** Drug/control ratios were calculated for all three sections. **(D)** The same drug/control ratios were used to construct concentration-inhibition plots, as it was explained in Fig. 1 (thick lines), which were then fitted by the Hill equation (thin lines). **(E)** The resulting IC50 (EIP) values are plotted against the same abscissae as in the B and C columns: conditioning pulse duration (**SDO**), interpulse interval duration (**RFI**), and membrane potential (**SSI**).

**Fig. 3.**
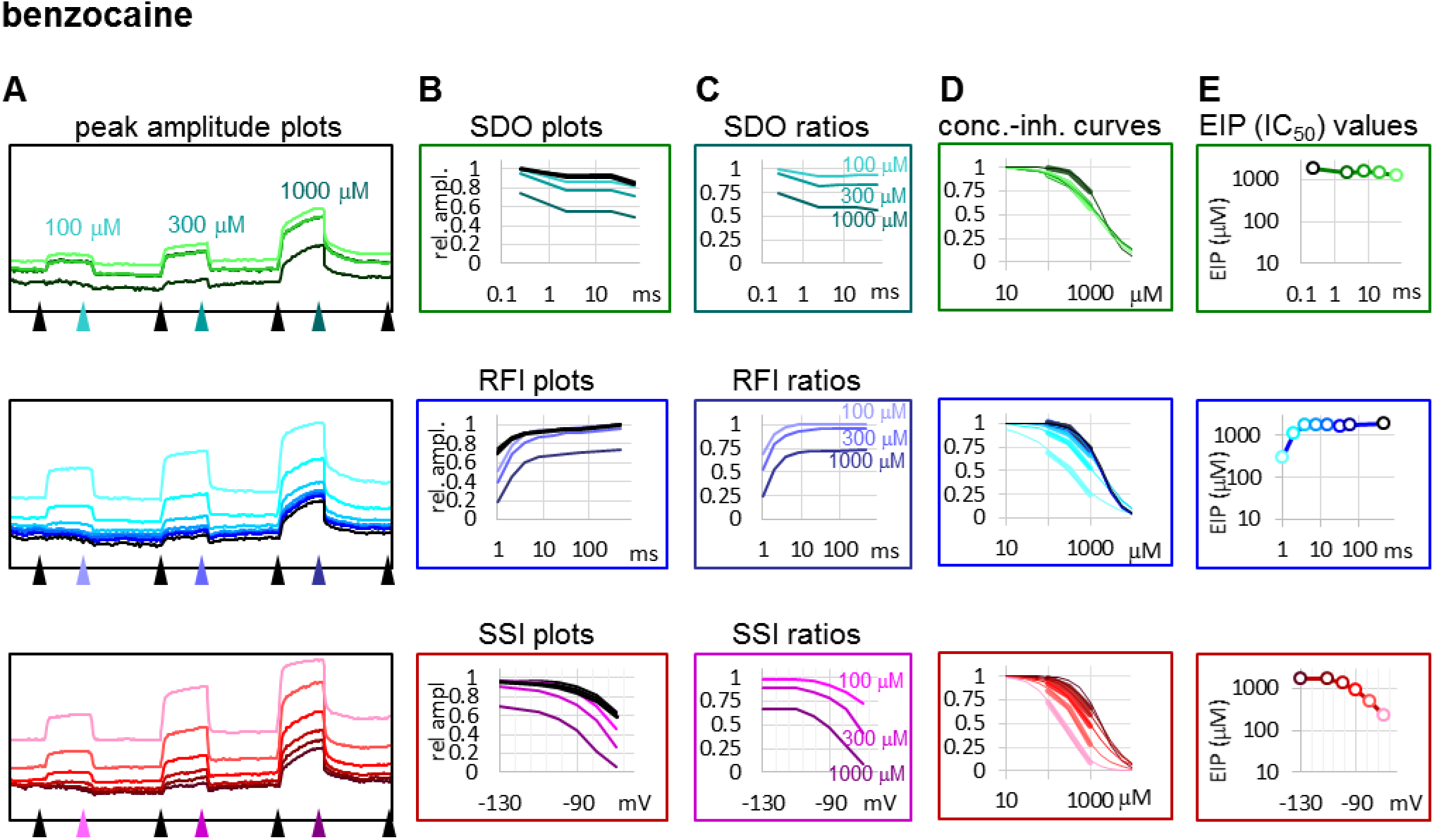
Construction of EIP plots for benzocaine. The figure illustrates how EIP plots are constructed on the example of a single measurement of the effect of three concentrations of benzocaine. **(A)** Peak amplitudes of currents evoked by the 17-pulse protocol were grouped into three sections: **SDO** (upper row), **RFI** (middle row), and **SSI** (lower row). **(B)** Amplitudes in control sections and during benzocaine perfusion were plotted against conditioning pulse duration (**SDO**), interpulse duration (**RFI**), and membrane potential (**SSI**). **(C)** Drug/control ratios were calculated for all three sections. **(D)** The same drug/control ratios were used to construct concentration-inhibition plots (thick lines), which were then fitted by the Hill equation (thin lines). **(E)** The resulting IC50 (EIP) values are plotted against the same abscissae as in the B and C columns.

**Fig. 4.**
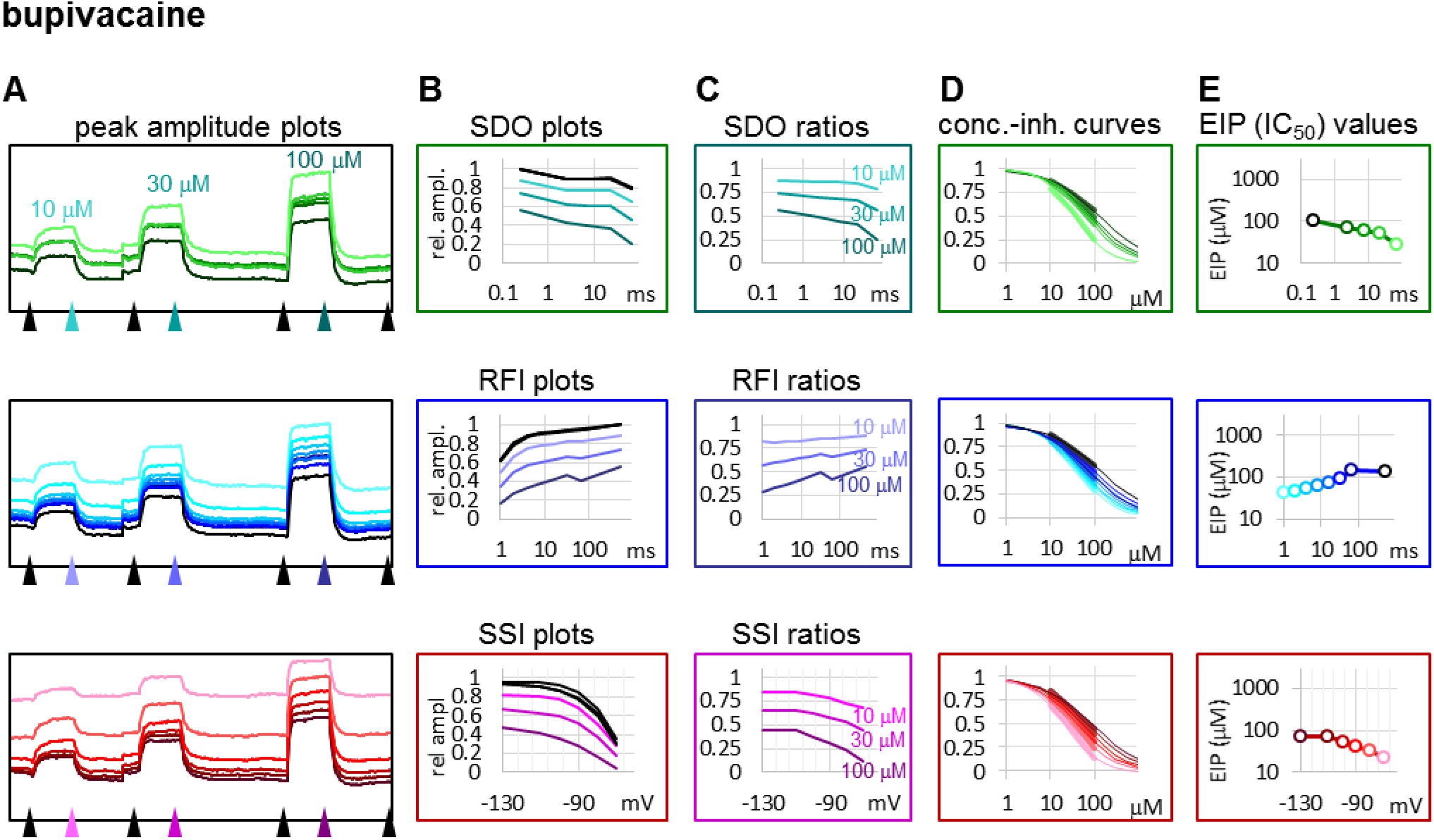
Construction of EIP plots for bupivacaine. The figure illustrates how EIP plots are constructed on the example of a single measurement of the effect of three concentrations of bupivacaine. **(A)** Peak amplitudes of currents evoked by the 17-pulse protocol were grouped into three sections: **SDO** (upper row), **RFI** (middle row), and **SSI** (lower row). **(B)** Amplitudes in control sections and during bupivacaine perfusion were plotted against conditioning pulse duration (**SDO**), interpulse duration (**RFI**), and membrane potential (**SSI**). **(C)** Drug/control ratios were calculated for all three sections. **(D)** The same drug/control ratios were used to construct concentration-inhibition plots (thick lines), which were then fitted by the Hill equation (thin lines). **(E)** The resulting IC50 (EIP) values are plotted against the same abscissae as in the B and C columns.

**Fig. 5.**
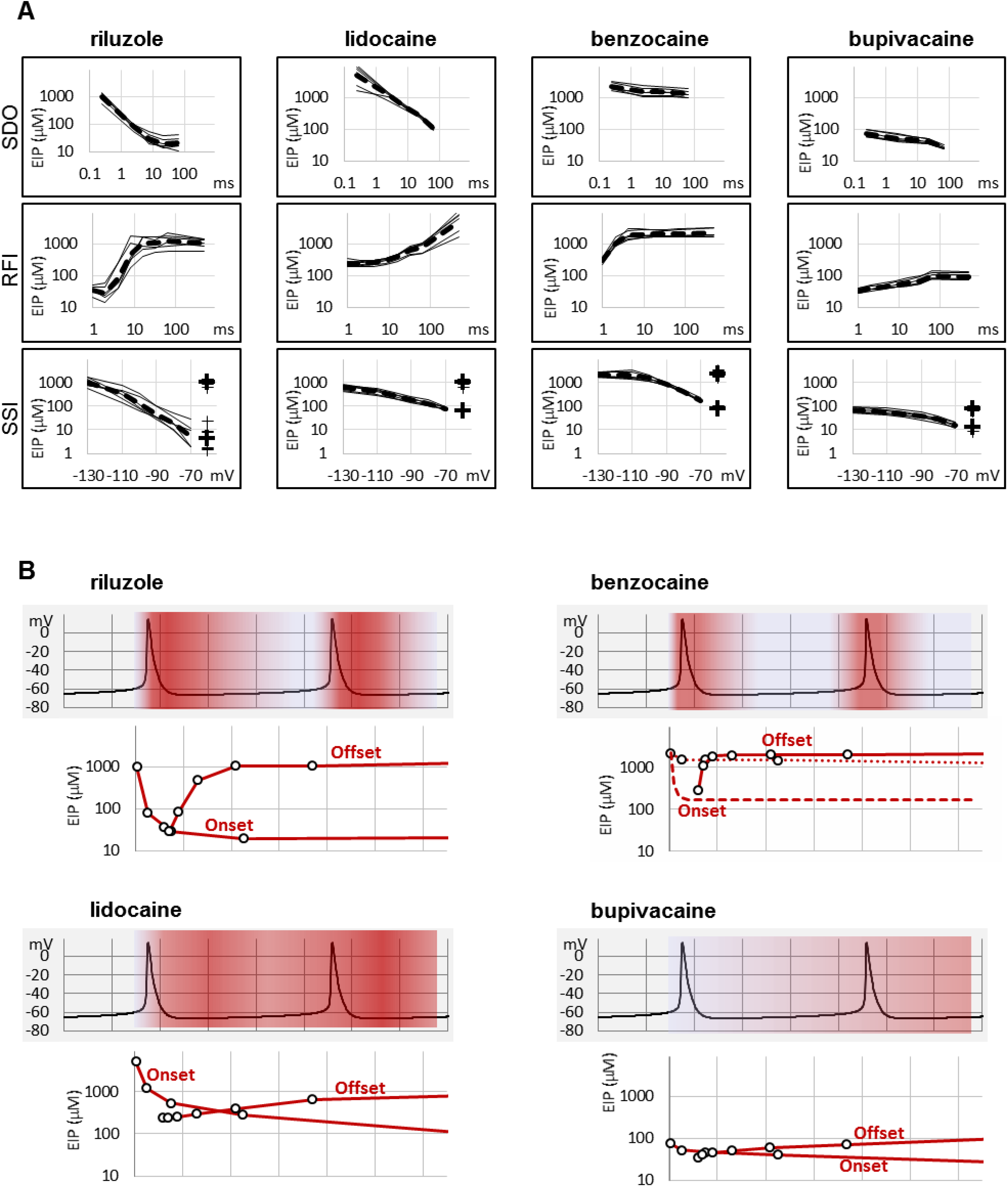
Summary of EIP plots for the four compounds, and the significance of micro-dynamics. **(A)** The micro-dynamics (EIP plots for **SDO** and **RFI**), and voltage-dependence of EIP are compared for n = 6 measurements for each of the four compounds. Thin lines show data from individual measurements, thick dashed lines show the geometric mean. For SSI data the calculated KR and KI values are also shown on the right side of the figures (thin crosses for individual measurements, and thick crosses for the geometric mean). **(B)** One example of how micro-dynamics is translated into frequency-dependent inhibition. The activity pattern of a simulated neuron is shown in the background, **SDO**-EIP and **RFI**-EIP plots are shown under the activity pattern, these are identical to the ones shown in dashed lines in panel A, upper and middle row, but on a linear time scale. Vertical grid: 10 ms. **SDO**-EIP plots were aligned with the upstroke of the first action potential because state-dependent onset is triggered by depolarization. **RFI**-EIP plots were aligned with the repolarization because hyperpolarization triggers recovery from inhibition. The intensity of red coloring illustrates how EIP is expected to change throughout the repetitive firing, and therefore what dynamics of inhibition is expected to be caused by the presence of the four compounds.

### Comparison of compound-specific properties of four different compounds

We chose three additional compounds in order to illustrate differences in micro-dynamics, even among closely related compounds, which showed similarly fast macro-dynamics. Lidocaine, benzocaine, and bupivacaine are all well-known local anesthetics. Examples of peak amplitude plots for all 17 pulses, **SDO**, **RFI**, and **SSI** plots, amplitude ratios, concentration-inhibition curves for all 17 pulses, and EIP plots are shown for all four compounds in Fig. 2 to 4. For each of the four compounds, we also calculated K_R_ and K_I_ estimates as described in Methods. Table 1 shows major parameters of inhibition, including K_R_ and K_I_ estimates, as well as K_R_/K_I_ ratios.

Lidocaine (Fig. 2) showed a slower micro-onset than riluzole. **SDO** ratio plots indicate that at 100 and 300 μM the onset did not reach its maximum even at the longest depolarization (64 ms) of the 17-pulse protocol. The recovery was also slower, substantial recovery only started after ~10 ms of hyperpolarization, and continued up to ~500 ms.

The fastest micro-dynamics we have encountered thus far was shown by benzocaine (Fig. 3). Recovery in fact was so fast that it compromised the measurement of **SDO**. At 2 ms gap duration, ~70 to 90 % of the channels have already recovered, therefore the 2.5 ms gap used in the **SDO** protocol was unable to reveal most of the inhibition caused by benzocaine binding. Although the plots of **SDO** ratios show the onset only on the remaining ~10 to 30% of channels, it is still discernible that the full onset has been completed within the shortest (2.5 ms) pulse. Calculation of K_I_ from SSI data (see Table 1) indicates that in fact more than half of the channels are inhibited by 100 μM of benzocaine, but this inhibition is difficult to detect whenever we use a hyperpolarizing gap before the test pulse.

When we tried to determine the kinetic behavior of bupivacaine (Fig. 4) we encountered the exact opposite of the problem with benzocaine: Even though the macro-dynamics was fairly fast (the time constant of macro-offset was 3.56 ± 0.43 ms), the micro-onset did not reach its maximum during the longest depolarization, and the micro-offset also could not reach equilibrium during the longest hyperpolarization. For this reason estimates of K_R_ and K_I_ are probably both incorrect, K_R_ was underestimated (*i.e.,* resting affinity would be less, if sufficient time was allowed for recovery) and K_I_ was overestimated (*i.e.,* inactivated affinity would be higher, if there was sufficient time for association).

In summary, we could detect radical differences between the micro-dynamics of four well-known sodium channel inhibitors. Summary of EIP values for the four compounds (n = 6 and their geometric mean for each one) are shown in Fig. 5A. On the EIP – SSI plots we also show K_R_ and K_I_ estimates for individual measurements (thin crosses), and their geometric mean (thick crosses). We propose that EIP values are the best means of characterizing micro-dynamics for individual compounds. In the Discussion, we examine why micro-dynamics is significant in predicting the effect of a compound on an active excitable cell. Table 1 shows all major concentration-independent, compound-specific parameters of inhibition, including K_R_ and K_I_ estimates, K_R_/K_I_ ratios, time constants of macro-offset (*τ_M-off_*), and time ranges of micro-onset and micro-offset. (Micro-onset and micro-offset could not be adequately described by exponentials, therefore instead we give the time range when the most substantial changes occur.)

## Discussion

### Micro-dynamics and frequency-selectivity

Riluzole has been originally described as an anti-epileptic compound^19^, and has been found to be an especially effective inhibitor at high firing frequencies^20–22^. In addition, it has been shown to selectively inhibit the persistent component (I_NaP_) of the sodium current^20^. We believe that both I_NaP_ selectivity and selective inhibition at high firing frequencies can be explained by its special micro-dynamics. We propose that preclinical assessment of sodium channel inhibitors should include evaluating their kinetic properties since these are important determining factors of their therapeutic potential.

To elucidate this, and to illustrate the significance of micro-dynamics, we chose four well-known compounds, with different micro-dynamics, and show their dynamically changing potency over a simulated firing of a simple, single-compartment neuron model (Fig. 5B). Our sole aim here is to illustrate how micro-dynamics of any compound can be interpreted, therefore we chose to visualize the EIP of the four compounds on the same firing pattern. We supposed that micro-onset is started upon depolarization, therefore the EIP plot from the **SDO** experiments was aligned with the upstroke of the action potential, and that micro-offset is started upon repolarization, therefore the EIP plot from the **RFI** experiments was aligned with the repolarization phase. The EIP plots are identical to the geometric mean curves shown in Fig. 5A, but here we use a linear time axis. The rates derived from non-physiological voltage patterns (square pulses between −130 and −10 mV) of course do not exactly match the rates under physiological membrane potential patterns, but they provide a rough estimate of the overall behavior of individual drugs. These estimates must be later verified by constructing models for their individual mechanisms of action, incorporating drug effects into the neuron model, and studying the interaction between different activity patterns and drug micro-dynamics.

In Fig.5B we show two subsequent action potentials of a cell that was induced to fire at ~25 Hz by a constant current injection. Changes in the intensity of red color illustrate dynamic changes in EIP during repeated action potentials. Dark red indicates high EIP (*i.e.*, IC_50_ approaches K_I_), white indicates low EIP (*i.e.*, IC_50_ approaches K_R_).

In the case of riluzole, we observed a massive state-dependence: with K_R_ = 1362 ± 205 μM and K_I_ = 5.63 ± 3.35 μM, the K_R_/K_I_ ratio was 242. The rate of micro-onset was somewhat slower than the action potential itself. This means that even if riluzole was present at a high concentration during an action potential, it would be ineffective: by the time riluzole could reach its maximal potency, the action potential would already be over. After the action potential riluzole stays very potent for ~10 ms, then it rapidly loses its potency again (for an explanation of this particularly fast micro-offset see Földi et al.^15^). This means that compounds with such micro-dynamics would be very potent inhibitors of high-frequency firing while being mostly ineffective at low frequencies. This is exactly what has been observed experimentally. For example, Desaphy et al.^21^ compared the frequency-dependent inhibition of seven compounds. Six of the compounds showed higher in vitro potency when the frequency of depolarizations was increased from 0.1 Hz to 10 Hz but did not change much between 10 and 50 Hz. Riluzole, on the other hand, only started to “realize its potential” above 10 Hz, the apparent affinity increased 48-fold, from IC50 = 43 μM (10 Hz) to IC50 = 0.9 μM (50 Hz). Inhibition of I_NaP_ may also contribute to inhibition of high-frequency burst firing^22^. As for I_NaP_ selectivity, we assume that it requires both a moderate micro-onset rate (not too fast so that it would miss the action potential, but not too slow so that it can inhibit I_NaP_ afterward) and a relatively fast offset rate (so that by the time of the next action potential it would lose its potency). While the potency of riluzole could both fully develop and fully fade away within 10 ms, changes in the EIP of lidocaine required more than 100 ms (for both the micro-onset and the micro-offset). This micro-dynamics, when plotted over the firing pattern (Fig. 5B), suggests that compounds with properties similar to lidocaine would follow firing frequencies with their micro-dynamics only up to ~5 Hz. Because micro-offset would require 100-200 ms, we would expect that for such compounds selective inhibition of the persistent component would best manifest itself in the 5-10 Hz range of firing frequencies. At higher frequencies, the extent of inhibition would simply depend on the average membrane potential in the course of firing activity.

Benzocaine is known as one of the fastest-acting sodium channel inhibitors, due to its small size and neutrality. Indeed, it is one of the few compounds, for which the whole process of partitioning, entry through the fenestration, and binding to the local anesthetic site has been observed in molecular dynamics simulations^23,24^. In our experiments we found extremely fast micro-offset kinetics, the offset was complete within 4 ms, and even the shortest interpulse interval (1 ms) already showed decreased potency (291 ± 22 μM, while the calculated K_I_ was 79.4 ± 3.9 μM). This was the reason, why the **SDO** protocol only detected a small fraction of the onset of drug action. Even though depolarization must have caused a definite increase in EIP, as we can see from the membrane potential dependence of EIP (Fig. 3), the 2.5 ms hyperpolarizing gap between pulses was enough to allow almost full recovery. Experimental results (which failed to detect the onset) are shown by open circles connected by a dotted line (Fig. 5B), while we suppose that the actual dynamics of micro-onset must have been complete within a few milliseconds (dashed line in Fig. 5B). It follows that significant frequency-dependence cannot be expected in the case of benzocaine, only at extremely high (>100 Hz) firing frequencies.

While benzocaine is a smaller, neutral compound that acts much faster than lidocaine, bupivacaine is larger, and has a higher pKa than lidocaine (therefore a somewhat larger fraction is charged at neutral pH). In an earlier comparative study, we found it to have higher potency, and slower onset/offset kinetics (only macro-dynamics was studied)^25^. In this study, we found that although its macro-dynamics was still relatively fast (macro-offset time constants were 1.72 ± 0.05 s for lidocaine, and 3.56 ± 0.43 s for bupivacaine, see Table 1), its micro-dynamics was slow, and therefore its EIP varied within a strikingly shallow range (between 27.3 ± 1.45 and 94.6 ± 10.7 μM). The K_R_/K_I_ ratio was only 7.48 (while for riluzole, lidocaine, and benzocaine, it was 242, 69.7, and 28.3, respectively). This can also be seen on the concentration-inhibition curves (Fig. 4), where the 17 different curves are very close to each other. The reason for a shallow micro-dynamics may be either that complete micro-onset of inhibition would require a depolarization even longer than 64 ms, or that the complete micro-offset would require a hyperpolarization even longer than 498 ms. Considering estimations of K_R_ and K_I_ from the literature (K_R_ ≈ 317.4 μM, and K_I_ ≈ 18.6 μM^26^; K_R_ ≈ 618.9 μM, and K_I_ ≈ 5.85 μM^25^), both could be the case for bupivacaine, therefore a more accurate assessment of its range of EIS values would require a protocol containing both longer depolarizations and longer hyperpolarizations. We presume that micro-onset and micro-offset both must be complete within 2-3 s since these processes cannot be slower than macro-offset, for which we observed a time constant of 3.56 s (Table 1). In the case of the simulated neuron firing at ~25 Hz, we suppose that development of inhibition would require several tens of action potentials, and cells would not substantially recover from inhibition between two action potentials unless the firing rate was less than 1 Hz.

### Micro-dynamics and persistent current selectivity

Micro-dynamics is a major determinant of the therapeutic profile. This is well known for the case of Class 1 antiarrhythmics, but the same principle can be applied to neuronal and skeletal muscle sodium channels, which can fire at a much higher rate. Micro-dynamics will determine which firing frequencies will be selectively inhibited, and it will also determine persistent current selectivity, as we have discussed above. Several compounds have been shown to selectively inhibit the persistent component of the sodium current (I_NaP_) over the transient component (I_NaT_) ^27–31^, riluzole being one of them^20^. The mechanism by which this selectivity is achieved, however, is not clear. It is not due to selectivity between sodium channel isoforms, but selectivity between conformations of the same channel isoforms may be part of the explanation. If an inhibitor compound has a higher affinity to the inactivated state, then I_NaT_ will be less affected because at the peak of the transient current there are few inactivated channels, but by the time I_NaT_ is over, and INaP is the only remaining sodium current, almost all channels have reached inactivated state. This is how INaP preference of riluzole was explained by Ptak *et al.* ^32^. However, almost all small molecule sodium channel inhibitors show higher affinity to inactivated state, and only a few of them are selective inhibitors of I_NaP_. We propose that – at least in the case of riluzole – micro-dynamics may be the key. Both micro-onset and micro-offset rates are crucial. Delayed micro-onset ensures that the transient component I_NaT_ is “missed” by the drug. Fast micro-offset, on the other hand, ensures, that upon hyperpolarization the inhibition is rapidly relieved, therefore I_NaT_ will be minimally affected at the next action potential. Obviously, there must be other possible mechanisms of I_NaP_ selectivity because this property is shared by compounds with much slower micro-dynamics, like e.g. ranolazine or phenytoin^27,29^. Details of these mechanisms still remain to be explored.

### Hill coefficients

We found that the best way to express dynamic changes in potency is by constructing concentration-inhibition curves for all depolarizing pulses and determining IC_50_ values. In principle, potency could be calculated from even a single concentration, if we supposed: i) one-to-one binding (i.e., the *n_H_ =* 1), and ii) that binding is equivalent with inhibition (channel block). In practice, however, for three out of the four compounds described here (riluzole, lidocaine, and benzocaine) we found Hill coefficients to vary widely, between ~0.5 and ~2 (Table 1, Fig. 1C, Figures 2–4). Steady-state availability data at different inhibitor concentrations can be converted to concentration-inhibition curves at different holding potentials (see Fig.1 A and B in Lenkey *et al.*^18^). In a study by Balser *et al.*^33^ this conversion was done on data with five different concentrations of lidocaine (see Fig. 6 and 7 in^33^), and the results gave *n_H_* values ranging from 0.64 to 1.83 (not given in the original paper, but could be reconstructed by fitting the data). Similarly to our results, *n_H_* < 1 values were observed at more negative holding potentials (−110 to −90 mV), where the potency was low (IC_50_ between 1900 and 2500 μM); while *n_H_* > 1 values were observed at less negative holding potentials (−70 to −50 mV). We assume, that *n_H_* < 1 values might reflect binding to multiple low-affinity binding sites which do not all cause full inhibition of conductance. Values greater than one, on the other hand, indicate positive cooperativity, which may come from multiple possible mechanisms, as has been discussed by Leuwer *et al.*^34^. They fitted V_1/2_ shift *vs.* concentration plots using a Hill-type exponent, which was found to range from 1.6 to2.1. These and our own data show that although *n_H_* may vary depending on the experimental protocol, it is quite common to find *n_H_* > 1 for sodium channel inhibitors, indicating that more than one inhibitor molecule is needed for effective inhibition, at least under certain experimental conditions. The binding of one molecule may induce or stabilize a conformation that is more favorable for binding of the second molecule. Alternatively, it is also possible that two or more bound molecules are required to effectively inhibit channels either by channel block or modulation. Even if we disregard modulation, channel block may in itself be more effective with two bound lidocaine molecules, as shown in a recent molecular dynamics simulation^35^.

## Conclusion

Therapeutic usefulness of sodium channel inhibitor drugs depends on their ability to selectively inhibit pathological activity of cells, which often manifests in hyperexcitability. Pathological hyperexcitability is involved in a number of disorders, including pain syndromes, epilepsies, muscle spasms, or arrhythmias. Each of the different types of hyperexcitability-related diseases has its characteristic dynamics of firing. To counteract them it is best to choose an inhibitor that has the precise dynamics that can selectively inhibit that certain pathological pattern of activity. We have demonstrated how much dynamic properties can differ even in the case of closely related compounds. Prediction of drug effects in an excitable tissue requires a thorough understanding of their mechanism of action, which includes the kinetic aspects of their state-dependent effects. In this study we aimed to derive compound-specific but concentration-independent descriptors of the mechanism of action, which can serve as a basis for building credible kinetic models for the simulation of drug-specific effects.

Thus far there has been no method available for a comparative study of drug onset/offset dynamics at a satisfactory throughput. Studies on the mechanism of action for individual drugs usually took several months to complete. Automated patch clamp instruments are capable of flawlessly performing complex experimental protocols, however, obtaining a large mass of complex information does not necessarily mean obtaining meaningful information. “Asking” the right question (by designing the right protocol) is not trivial, and neither is deciphering a relevant “answer” from a huge amount of data. Furthermore, one needs to find the right degree of automation, which allows fast analysis, but also allows manual handling and monitoring of data. We assume that the method described in this study and its prequel will inspire other groups to use automated patch clamp instruments more creatively, in order to better exploit their potential.

## List of non-standard abbreviations

SDO: “state-dependent onset” protocol
RFI: “recovery from inactivation” protocol
SSI: “steady-state inactivation” protocol
EIP: effective inhibitor potency

